# *CcLBD25* functions as a key regulator of haustorium development in *Cuscuta campestris*

**DOI:** 10.1101/2021.01.04.425251

**Authors:** Min-Yao Jhu, Yasunori Ichihashi, Moran Farhi, Caitlin Wong, Neelima R. Sinha

## Abstract

Parasitic plants reduce yield of crops worldwide. *Cuscuta campestris* is a stem parasite that attaches to its host, using haustoria to extract nutrients and water. We analyzed the transcriptome of six *C. campestris* tissues and identified a key gene, *CcLBD25*, as highly expressed in prehaustoria and haustoria. Our gene co-expression networks (GCN) from different tissue types and laser-capture microdissection (LCM) RNA-Seq data indicate that *CcLBD25* could be essential for regulating cell wall loosening and organogenesis. We employed host-induced gene silencing (HIGS) by generating transgenic tomato hosts that express hairpin RNAs to target and down-regulate *CcLBD25* in the parasite. Our results showed that *C. campestris* growing on *CcLBD25* RNAi transgenic tomatoes transited to the flowering stage earlier and had less biomass compared with *C. campestris* growing on wild type host. This suggests that the parasites growing on the transgenic plants were stressed due to insufficient nutrient acquisition. Anatomy of *C. campestris* haustoria growing on *CcLBD25* RNAi tomatoes showed reduced pectin digestion and lack of searching hyphae, which interfered with haustorium penetration and the formation of vascular connections. We developed an *in vitro* haustorium (IVH) system to assay the number of prehaustoria produced on strands from *C. campestris*. When *C. campestris* was grown on *CcLBD25* RNAi tomatoes or wild type tomatoes, the former produce fewer prehaustoria than the latter, indicating that down-regulating *CcLBD25* may affect haustorium initiation. The results of this study shed light on the role of *CcLBD25* in haustorium development and might help to develop a parasite-resistant system in crops.

**One-sentence summary:** *CcLBD25* plays a pivotal role in haustorium initiation, regulating pectin digestion, and searching hyphae development during the haustorium penetration process.

## Introduction

Parasitic plants are heterotrophic, reducing the yields of crops worldwide (Agrios, 2005; Yoder and Scholes, 2010). They parasitize host plants using specialized organs known as haustoria, which extract nutrients and water from the hosts. *Cuscuta* species (dodders) are stem holoparasites without functional roots and leaves. Stems of *Cuscuta* spp. coil counterclockwise around their host and then form a series of haustoria along the stems to attach to their hosts (Furuhashi et al., 2011; Alakonya et al., 2012). *Cuscuta campestris* is one of the most widely distributed and destructive parasitic weeds. A better understanding of the underlying molecular mechanisms of *C. campestris* haustorium development will aid in the application of parasitic weed control and producing parasitic plant-resistant crops.

Many previous studies have identified the key factors needed for seed germination, host recognition, and haustorium induction and growth in root parasites (Shen et al., 2006; López-Ráez et al., 2009; Yoder and Scholes, 2010). Focusing on haustorium development, a previous study indicated that root parasitic plants co-opted the mechanism of lateral root formation in haustorium organogenesis (Ichihashi et al., 2017). LATERAL ORGAN BOUNDARIES DOMAIN (LBD) family of transcription factors (TFs) are reported to be crucial in both lateral root formation in non-parasitic plants and haustorium developmental programming in root parasites (Ichihashi et al., 2020). In non-parasitic model plants, like Arabidopsis, LBD genes are shown to be involved in auxin signaling, interact with AUXIN RESPONSE FACTORs (ARFs) and promote lateral root formation (Mangeon et al., 2010; Porco et al., 2016). Further, LBD orthologs are reported to be upregulated during the haustorium development stage at attachment sites in root parasitic plants like *Thesium chinense* (Ichihashi et al., 2017) and *Striga hermonthica* (Yoshida et al., 2019). On the other hand, the molecular pathways regulating haustorium development in stem parasitic plants are still largely unexplored. Although a few gene orthologs that regulate auxin accumulation during lateral root development in non-parasitic plants are found to be expressed in *Cuscuta* seedling and stems, whether these genes are also involved haustorium formation is still unknown (Ranjan et al., 2014). Our previous studies showed that the *SHOOT MERISTEMLESS*-like (*STM*) plays a role in *Cuscuta* spp. haustorium development (Alakonya et al., 2012). These results suggest that *Cuscuta* spp. might have repurposed the shoot developmental programs into haustorium organogenesis, but whether *Cuscuta* spp. also co-opted the lateral root programming system into haustorium development remains an open question.

In this study, we provide an insight into the gene regulatory mechanisms of haustorium organogenesis and identify one of the LBD transcription factors, *CcLBD25*, as a vital regulator of *C. campestris* haustorium development. This discovery supports the hypothesis that stem parasitic plants adapted both shoot and root molecular machinery into haustorium formation. Using detailed transcriptome analysis and gene coexpression networks coupled with cellular and developmental phenotypic assays, we also show that *CcLBD25* is not only involved in haustorium initiation through the auxin signaling, but also participates in other aspects of haustorial developmental reprogramming, including cell wall loosening, searching hyphae development, and other phytohormone mediated signaling pathways. The results of this study will not only shed light on the field of haustorium development in stem parasitic plants but also help develop a potential universal parasitic weed-resistant system in crops to reduce economic losses caused by both root and stem parasites.

## Results

### Establishing genomic resources for *C. campestris* and constructing gene coexpression networks regulating haustorium formation

In this study, we analyzed the transcriptome of different *C. campestris* tissues, including seeds, seedlings, stems, prehaustoria, haustoria, and flowers, grown on the tomato (*Solanum lycoperscum*) Heinz 1706 (H1706) cultivar and *Nicotiana benthamiana* (*N. benthamiana*) (Ranjan et al., 2014) by mapping to the recently available genome of *C. campestris* (Vogel et al., 2018). In general, seed tissues have distinctively different gene expression profiles compared to all other tissues (Supplemental Fig. 1). In addition, the expression patterns in invasive tissues (prehaustoria and haustoria) and non-invasive tissues are also disparate (Supplemental Fig. 1). We conducted PCA analysis and noticed the genes that are highly expressed in invasive tissues can be separated from the genes that are highly expressed in non-invasive tissue on PC1 (Fig. 1A, B). To identify the genes that might be involved in haustorium development, we performed clustering analysis using self-organizing maps (SOM) in R and identified a cluster enriched with genes that are highly expressed in both prehaustoria and haustoria tissues (SOM9 - Fig. 1, Supplemental Fig. 2 and 3).

**Figure 1.**
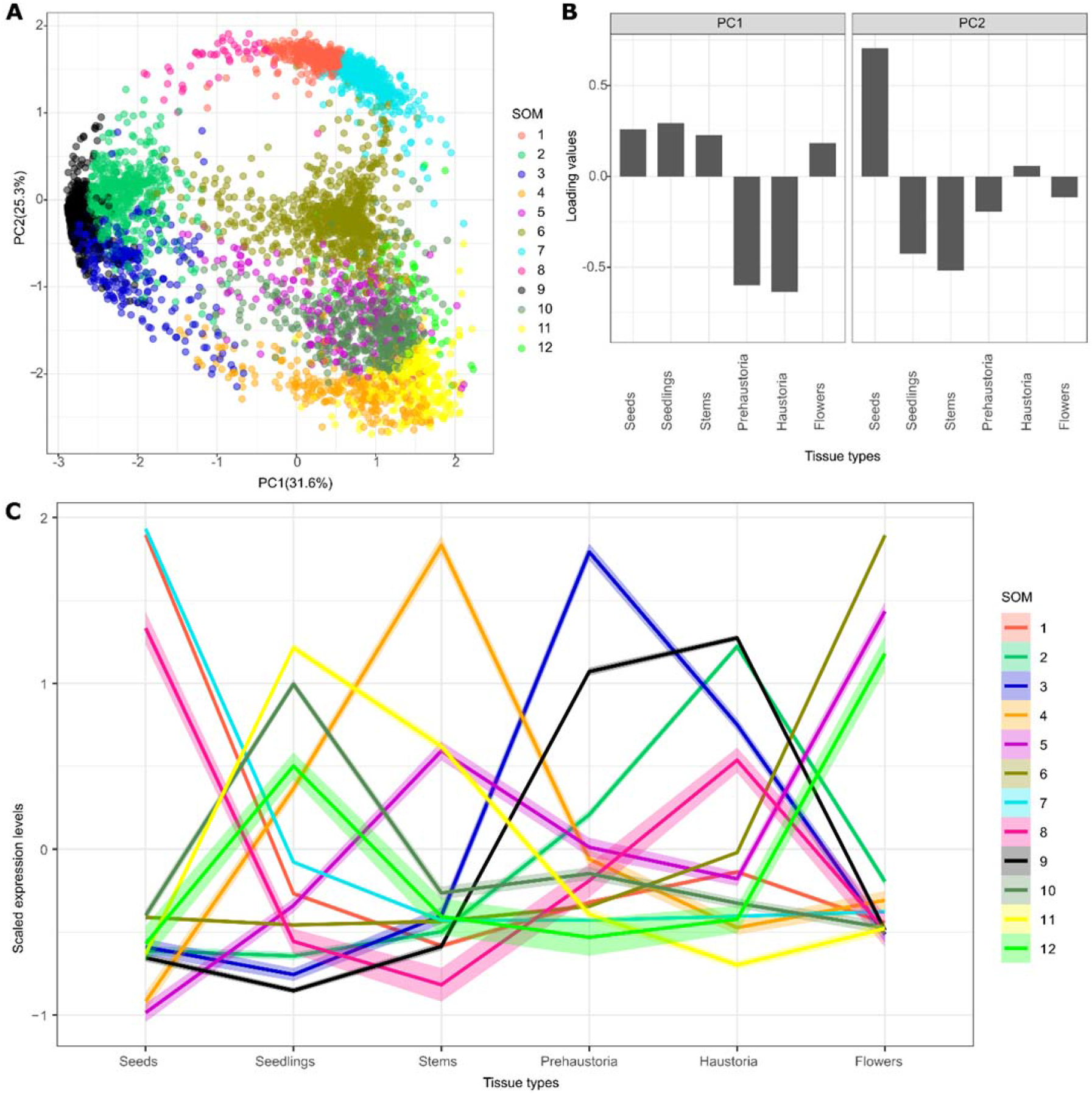
PCA analysis with SOM clustering of gene expression in *C. campestris* tissue type RNA-Seq data mapped to *C. campestris* genome. (A) PCA analysis based on gene expression across different *C. campestris* tissues. Each dot represents a gene and is in the color indicating their corresponding SOM group. (B) Loading values of PC1 and PC2. PC1 separates the genes that are specifically expressed in intrusive tissues (prehaustoria and haustoria) from those that are expressed in non-intrusive tissues. PC2 divides the seed-specific gene from other genes. (C) Scaled expression levels of each SOM group across different *C. campestris* tissue types. Each line is colored based on corresponding SOM groups.

We focused on the genes contained in this SOM9 cluster and constructed a gene coexpression network (GCN). Using the fast greedy modularity optimization algorithm to analyze the GCN community structure (Clauset et al., 2004) and visualizing the network using Cytoscape (Cline et al., 2007), we noticed this SOM9 GCN is composed of three major modules (Fig. 2A). Since the current gene annotation of *C. campestris* genome is not as complete as that of most model organisms, we used BLAST to combine our previously annotated transcriptome with current *C. campestris* genome gene IDs (Supplementary Table 1). With this more comprehensive annotation profile, we further conducted GO enrichment analysis using the TAIR ID for each *C. campestris* gene in the network to identify the major GO term for each module. Based on our GO enrichment results, the major biological process of module 1 can be classified as “plant-type cell wall loosening” and the cellular component of module 1 is “extracellular region and intracellular membrane-bounded organelle” (Fig. 2A, B). This result indicates the genes contained in module 1 are mostly involved in cell wall loosening, which is needed for the haustorium to penetrate through the host tissue. On the other hand, the major biological processes of module 3 are “transport, response to hormones, secondary metabolite biosynthetic process, and regulation of lignin biosynthetic process”. The molecular function of module 3 is “transmembrane transporter activity” and the cellular component of module 3 is “plasma membrane” (Fig. 2A). This analysis suggests that these genes might be involved in later stages of development and nutrient transport from the host to the parasite once a connection is established between the host and the parasite.

**Figure 2.**
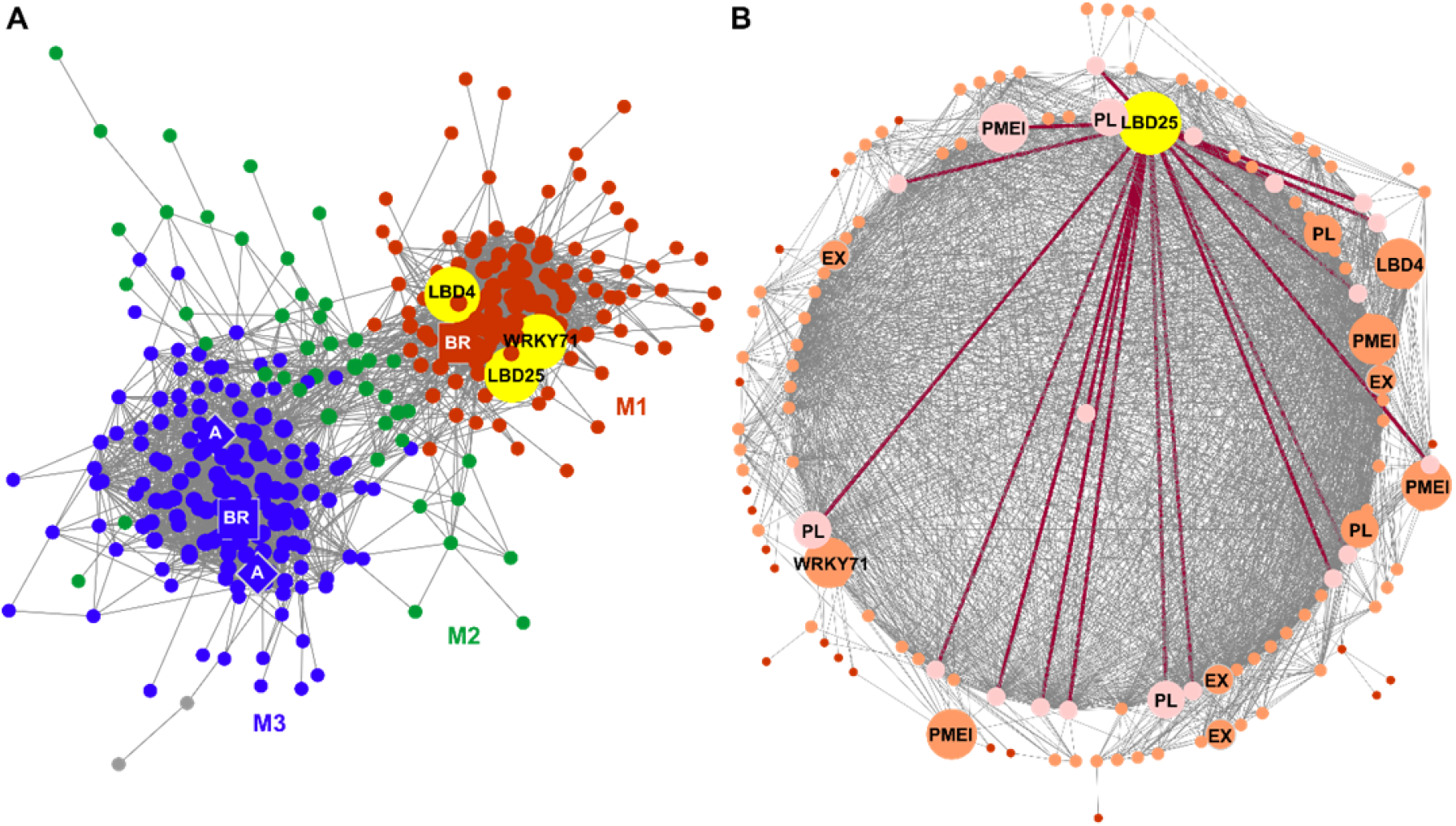
SOM9 gene-coexpression networks (GCNs) from *C. campestris* tissue type RNA-Seq data. (A) GCN of genes that are classified in SOM9, which includes genes that are highly expressed in both prehaustoria and haustoria. This SOM9 GCN is composed of three major modules. Red indicates genes in Module 1. Green indicates genes in Module 2. Blue indicates genes in Module 3. The only three transcription factors (TFs) in module 1 are labeled in yellow. (B) GCN of genes that are classified in SOM9 Module 1. Dark red lines indicate the connection between *CcLBD25* and its first layer of neighbors. The genes that are first layer neighbors of *CcLBD25* are labeled in pink. The genes that are second layer neighbors of *CcLBD25* are labeled in orange. The only three TFs and cell wall loosening related gene are highlighted and labeled in the network. (A, B) PL, pectin lyase. PMEI, pectin methyl-esterase inhibitor. EX, expansin. A, auxin efflux carrier-like protein. BR, brassinosteroid insensitive 1-associated receptor kinase 1-like.

To identify the key regulators in the haustorium penetration process, we focused on genes in module 1 and calculated the degree centrality and betweenness centrality scores of each gene within this group. Many central hub genes in module 1 are proteins or enzymes involved in cell wall modifications, like pectin lyases, pectinesterase inhibitors, and expansins (Fig. 2B). To find the upstream regulators of these pathways, we focused on transcription factors that are classified in module 1. Intriguingly, only three transcription factors are included in module 1: *CcLBD25* (Lateral Organ Boundaries Domain gene 25), *CcLBD4*, and *CcWRKY71* (Fig. 2B). According to the gene coexpression network of SOM9, we noticed that *CcLBD25, CcLBD4*, and *CcWRKY71* share several common first layer of neighbors (Figure 2). Based on previous reports, *AtLBD25* regulates lateral root development in *Arabidopsis* by promoting auxin signaling (Dean et al., 2004; Mangeon et al., 2010). Furthermore, an *LBD25* orthologue (*TcLBD25*) in *Thesium chinense*, a root parasitic plant in the Santalaceae family, was also detected to be upregulated during the haustorium development process (Ichihashi et al., 2017). With these serendipitous pieces of evidence, we suspected that *CcLBD25* may regulate haustorium formation and the parasitism process in *C. campestris*.

To understand the role of *CcLBD25* and the potential connection with other genes, we included genes in SOM2 (genes that are only highly expressed in haustoria) and SOM3 (genes that are only highly expressed in prehaustoria) to build a more comprehensive GCN (Fig. 1, Supplemental Fig. 2 and 3). Based on the community structure analysis, this comprehensive network is composed of three major modules (Fig. 3A). Based on our GO enrichment results, the major biological process of module 3 is plant-type cell wall loosening and the major biological processes of module 1 are morphogenesis of a branching structure, plant organ formation, and several hormone responses and biosynthetic processes. *CcLBD25* itself is placed in module 1, but *CcLBD25* has many first layer connection with genes that are classified in module 1 or module 3 (Fig. 3A, C). This result indicates that *CcLBD25* might play a role in connecting genes involved in different pathways or aspects of haustorium development. Furthermore, by coloring the network with their corresponding SOM groups, we noticed that even though *CcLBD25* itself is in SOM9, many of the *CcLBD25* first and second layers of neighbors are in SOM2 and SOM3 (Fig. 3B, D). Thus, *CcLBD25* might be a key regulator of the haustorium development process in both early and late stages of haustorium development, and may also play a critical role in coordinating the function of genes that are only expressed in discrete developmental stages.

**Figure 3.**
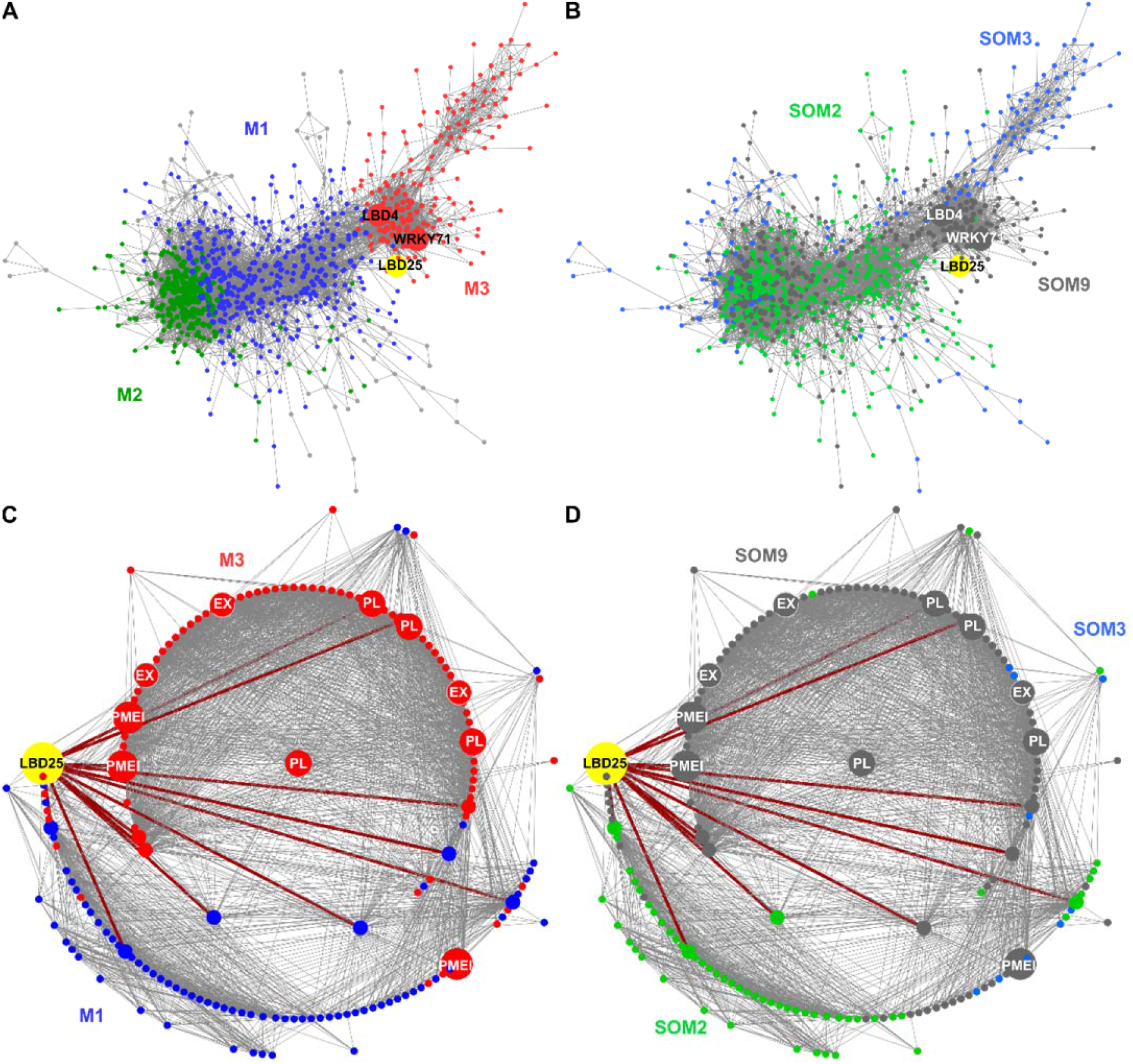
GCNs of SOM2, 3, 9 genes based on *C. campestris* tissue type RNA-Seq data. (A) GCN of genes that are in SOM2, SOM3, and SOM9 with colors based on network modules. This SOM2+SOM3+SOM9 GCN is composed of three major modules. Red indicates genes in Module 1. Green indicates genes in Module 2. Blue indicates genes in Module 3. (B) GCN of genes that are in SOM2, SOM3, and SOM9 with colors based on SOM clustering groups. Green indicates genes in SOM2. Blue indicates genes in SOM3. Grey indicates genes in SOM9. SOM2 includes genes that are only highly expressed in haustoria and SOM3 includes genes that are only highly expressed in prehaustoria. (C) GCN of *CcLBD25* and its first and second layer of neighbors with colors based on network modules. (D) GCN of *CcLBD25* and its first and second layer of neighbors with colors based on SOM clustering groups. (C, D) Red lines indicate the connection between *CcLBD25* and its first layer of neighbors. PL, pectin lyase. PMEI, pectin methyl-esterase inhibitor. EX, expansin.

### Zooming into tissue specific expression using laser-capture microdissection (LCM) coupled with RNA-seq

Our first transcriptome data came from hand collected tissue samples. To further dissect *Cuscuta* haustorium developmental stages, we used laser-capture microdissection (LCM) with RNA-seq to analyze only pure haustorial tissues from three different haustorium developmental stages (Supplemental Fig. 4). Based on a previous study, changes in the levels of jasmonic acid (JA) and salicylic acid (SA) are observed about 36-48 hours after first haustorial swelling, which is about 4 days post attachment (DPA) (Runyon et al., 2010). We also noticed that haustorium growth is a continuous process on *C. campestris*, so all developmental stages of haustoria can be found on the same strand at the 4 DPA time point. Therefore, we focused on 4 DPA and defined three developmental stages based on their haustorium structure: early (the haustorium has just contacted the host), intermediate (the haustorium has developed searching hyphae but has not formed vascular connections), and mature (a mature haustorium with continuous vasculature between host and parasite) (Supplemental Fig. 4). *C. campestris* haustorium tissues were collected using LCM at these three developmental stages from *C. campestris* attached on H1706 and subjected to RNA-Seq (Supplemental Fig. 4).

Next, we mapped our LCM RNA-Seq data to the *C. campestris* genome. Visualizing the gene expression changes using multidimensional scaling showed that the expression profile of the mature-stage is distinct from the early and intermediate stages (Supplemental Fig. 5). We then conducted clustering analyses using SOM to group genes based on their expression patterns at these three different developmental stages (Fig. 4). According to our PCA analysis, PC1 obviously separated genes that are specifically expressed in the mature-stage from those expressed in the other two stages, and PC2 distinguished the genes expressed in early stage from intermediate stage (Supplemental Fig. 6). Interestingly, and similar to what was seen in our tissue type transcriptome data, *CcLBD25* is grouped in SOM6, which is the cluster of genes that are relatively highly expressed in both early-stage and mature-stage (Fig. 4A, B, and Supplemental Fig. 7). To investigate gene regulatory dynamics within the haustorium developmental process, we used the same gene list from tissue type RNA-Seq SOM9 and constructed a new GCN of these genes based on the LCM RNA-Seq expression profiles (Fig. 4C). By using the same gene list, but the expression dataset from samples of precisely collected haustorial cells, we obtained more detailed regulatory connections between genes by comparing the tissue type GCN and LCM GCN (Fig. 2A, 4C). Based on the fast greedy community structure analysis, this LCM GCN is composed of three major modules and *CcLBD25* is in module 1 (Fig. 4C). According to our GO enrichment results, the major biological process for module 1 is plant-type cell wall loosening and for module 3 is brassinosteroid mediated signaling pathway (Fig. 4C). In addition to cell wall loosening related enzyme encoding genes forming central hubs, we noticed *CcLBD25* is the TF with the highest connection in module 1 and has many connections with cell wall loosening-related genes (Fig. 4C, D). Zooming in to focus on *CcLBD25*, we noticed that the *CcLBD25* first and second layers of neighbors are genes classified in module 1 or module 3, indicating that *CcLBD25* might play a role in connecting these two pathways. Many of the *CcLBD25* first and second layers of neighbors are pectin degradation related genes, like *PL* and *PMEI*. On the other hand, *CcLBD4* is not in the LCM GCN and *CcWRKY71* is at a marginal location with only one connection. This result provided further support for our hypothesis that *CcLBD25* is the major TF regulating cell wall modification in the haustorium penetration process. *CcLBD4* and *CcWRKY71* might also be key regulators, but are likely involved in a different aspect of haustorium development. Thus, we focused our attention on understanding the function of *CcLBD25* in haustorium development.

**Figure 4.**
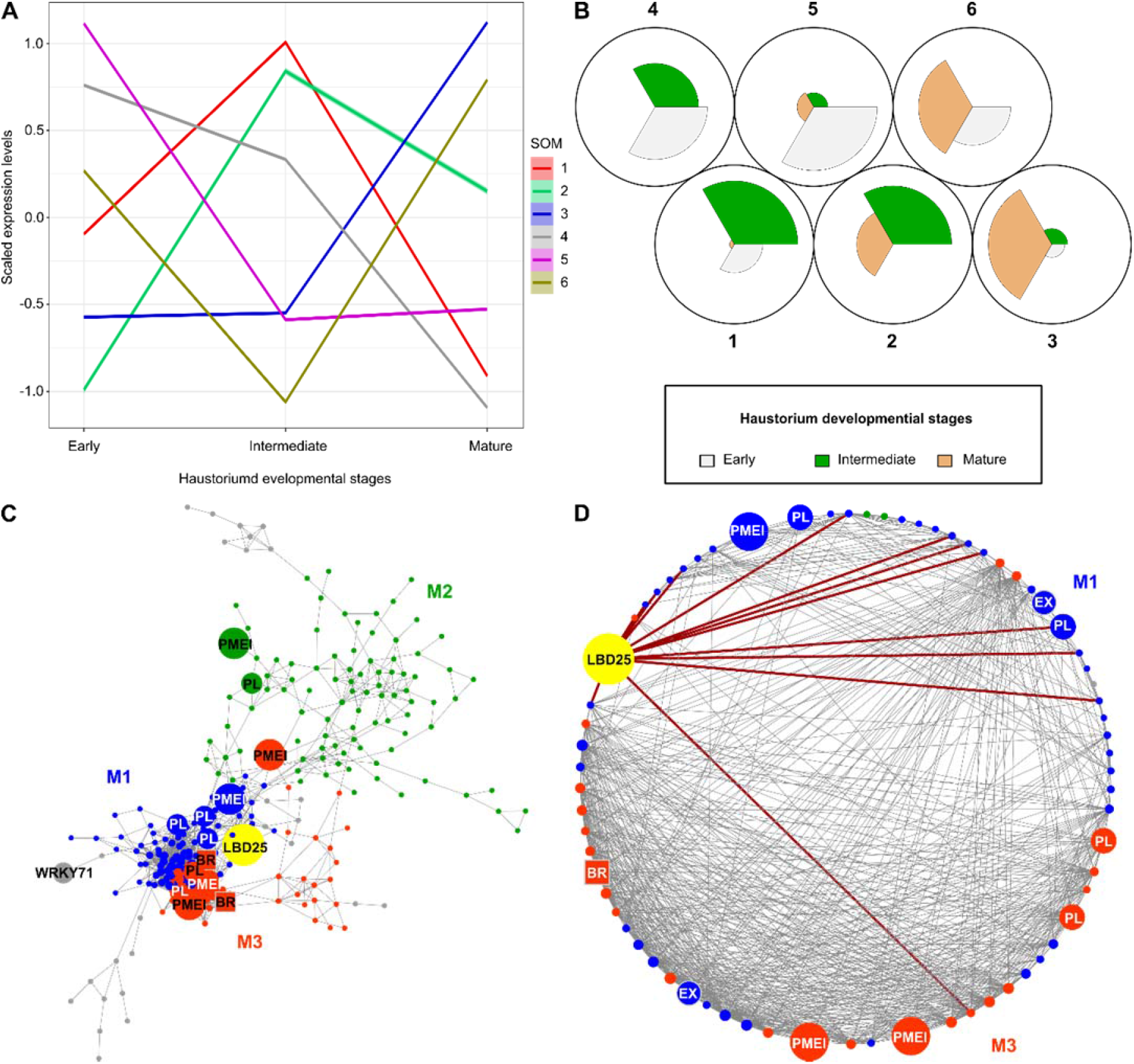
SOM clustering and GCNs of gene expression in *C. campestris* haustoria across three developmental stages in LCM RNA-Seq data. (A) Scaled expression levels of each SOM group across three haustorium developmental stages. Each line is colored based on corresponding SOM groups. (B) A code plot of SOM clustering illustrating which developmental stages are highly represented in each SOM group. Each sector represents a developmental stage and is in the color indicating their corresponding developmental stage. (C) GCN based on LCM RNA-Seq expression profiles with genes in tissue type RNA-Seq SOM9. Blue indicates genes in Module 1. Green indicates genes in Module 2. Red indicates genes in Module 3. Pectin degradation related gene are highlighted and labeled in the network. (D) GCN of *CcLBD25* and its first and second layer of neighbors with colors based on network modules. Dark red lines indicate the connection between *CcLBD25* and its first layer of neighbors. (C-D) PL, pectin lyase. PMEI, pectin methyl-esterase inhibitor. EX, expansin. BR, brassinosteroid insensitive 1-associated receptor kinase 1-like.

### Cross-species RNAi (Host-Induced Gene Silencing) *CcLBD25* effects whole-plant phenotypes and reduces the parasite fitness

In our previous studies, we found cross-species transport of mRNAs and siRNAs between *C. campestris* and their hosts, and demonstrated host-induced gene silencing (HIGS) (Runo et al., 2011; Alakonya et al., 2012). Many previous studies have also shown that large-scale mRNA and small RNAs are transported through the haustorium connections in *Cuscuta* species (Kim et al., 2014; Johnson et al., 2019). Therefore, we generated transgenic host tomato with hairpin RNAs that target and down-regulate *CcLBD25* after the parasite forms the first attachment and takes up RNAs from the host (Supplemental Fig. 8). When *C. campestris* grows on wild-type tomato hosts, *CcLBD25* is highly expressed in invasive tissues (Fig. 5A, B). However, *CcLBD25* expression levels are significantly knocked-down in the tissues on and near the attachment sites of *C. campestris* plants that are growing on *CcLBD25* RNAi transgenic plants (Fig. 5B). If *CcLBD25* is important in haustorium development and parasitism, then down-regulating *CcLBD25* should influence haustorium structure or formation and might also affect nutrient transport. To verify our hypothesis, we measured flowering time in *C. campestris* growing on various tomato hosts. The result showed that parasites growing on *CcLBD25* RNAi transgenic tomatoes transitioned to the flowering stage and subsequently senesced earlier than those growing on wild types (Fig. 5C). Based on previous studies, many plant species respond to environmental stress factors by inducing flowering (Wada and Takeno, 2010; Riboni et al., 2014). This early transition to the reproductive stage and senescence in *C. campestris* grown on *CcLBD25* RNAi plants suggests that *C. campestris* was growing under stress likely because of nutrient deficiency.

**Figure 5.**
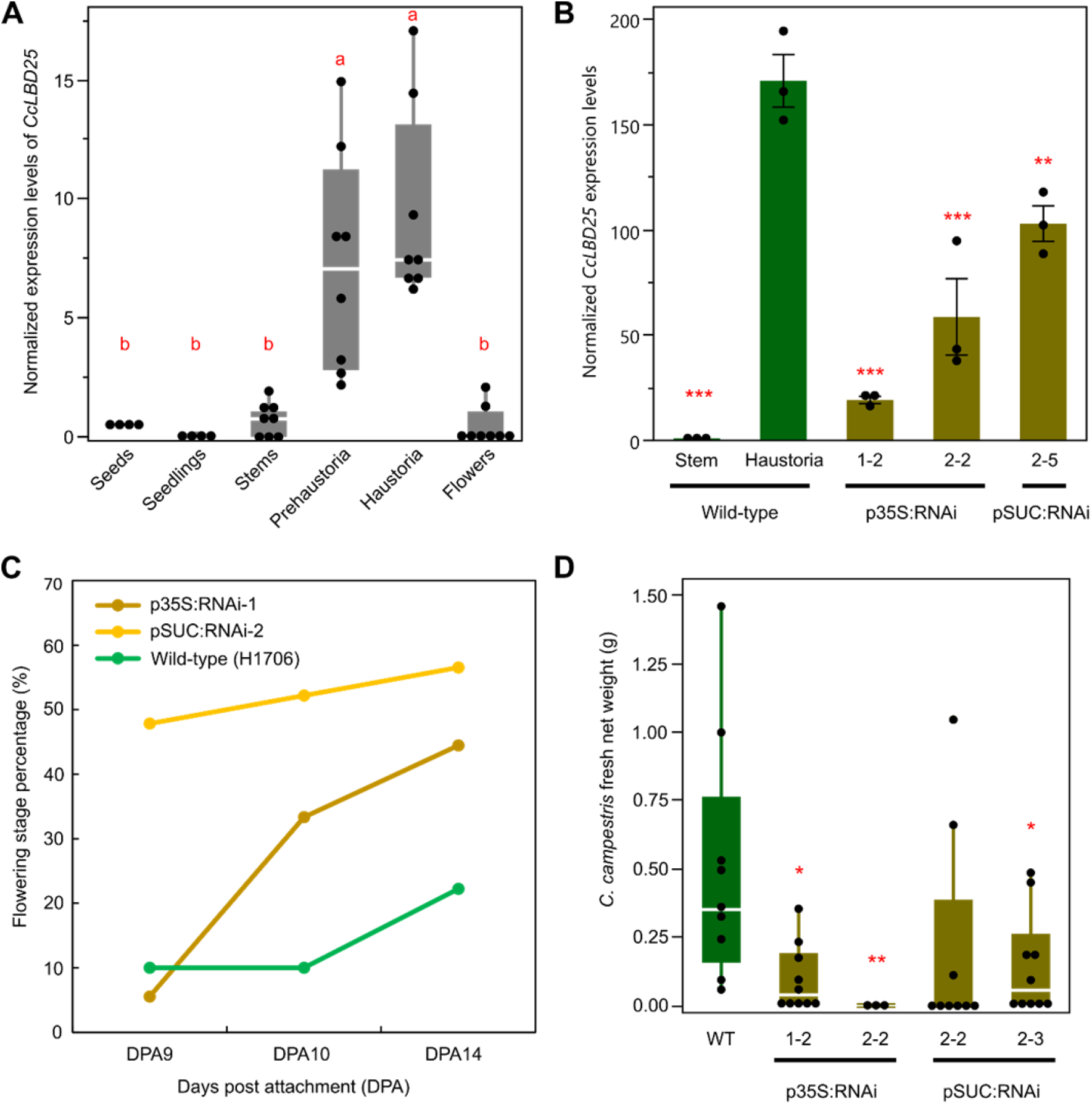
Gene expression levels and whole-plant phenotypes of *C. campestris* growing on Host-Induced Gene Silencing (HIGS) *CcLBD25* RNAi transgenic plants. (A) The normalized expression level of *CcLBD25* in six different tissue types of *C. campestris* from RNA-Seq data. Data presented are assessed using pair-wise comparisons with the Tukey test. P-value of the contrasts between “a” and “b” are less than 0.001. (B) Expression levels of *CcLBD25* in *C. campestris* haustoria grown on wild-type tomatoes and *CcLBD25* RNAi transgenic plants. Data presented are assessed using one-tailed Welch’s t-test with wild-type haustoria as control. “*” p-value < 0.05. “**” p-value < 0.01. “***” p-value < 0.005. (C) The flowering time of *C. campestris* growing on wild-type tomatoes and *CcLBD25* RNAi transgenic plants. The early transition to the flowering stage indicates that *C. campestris* may be growing under stress conditions because they might not obtain sufficient nutrients from their host. DPA, day post attachment. (D) Biomass of *C. campestris* growing on wild-type tomatoes and *CcLBD25* RNAi transgenic plants. Fresh net weights of *C. campestris* were measured in gram (g). Data presented are assessed using one-tailed Welch’s t-test with wild-type (WT) as control. “*” p-value < 0.05. “**” p-value < 0.01. (B, C, D) p35S:RNAi indicates the transgenic plants with the 35S promoter driving *CcLBD25* RNAi construct. pSUC:RNAi indicates the transgenic plants with the SUC2 promoter driving *CcLBD25* RNAi construct.

To verify if down-regulating *CcLBD25* affects the ability of the parasite to acquire resources from the host, we also measured the biomass of *C. campestris* grown on wild-type H1706 and *CcLBD25* RNAi transgenic plants. At 14 days post attachment (DPA), we noticed that *C. campestris* plants grown on *CcLBD25* RNAi transgenic tomatoes have less biomass compared with the *C. campestris* plants grown on wild-type H1706 (Fig. 5D). Both whole-plant level phenotypes suggest that *CcLBD25* might be involved in haustorium development and knocking down the expression level of *CcLBD25* influences the ability of *C. campestris* to establish connections with hosts and interferes with parasite nutrient acquisition.

### Down-regulation of *CcLBD25* leads to structural changes in haustoria

To verify the crucial role *CcLBD25* plays in haustorium development and to investigate how down-regulating *CcLBD25* affects haustorium structure and the parasitism process, we prepared 100 µm thick fresh haustorium sections using a vibratome and stained them with Toluidine Blue O (O’Brien et al., 1964). In wild-type haustorium sections, we could observe searching hyphae penetrate the host cortex region and transform into xylic or phloic hyphae as they connected to host xylem and phloem (Fig. 6A, C, E). However, we observed that many haustoria growing on *CcLBD25* RNAi transgenic tomatoes form a dome shape structure and lack searching hyphae (Fig. 6B, D, F, and Supplemental Fig. 9). This result indicates that *CcLBD25* might be involved in searching hyphae development. Therefore, knocking down of *CcLBD25* affects the ability of *C. campestris* to establish connections with the host vascular system and leads to nutrient deficiency as observed in the whole-plant level phenotypes.

**Figure 6.**
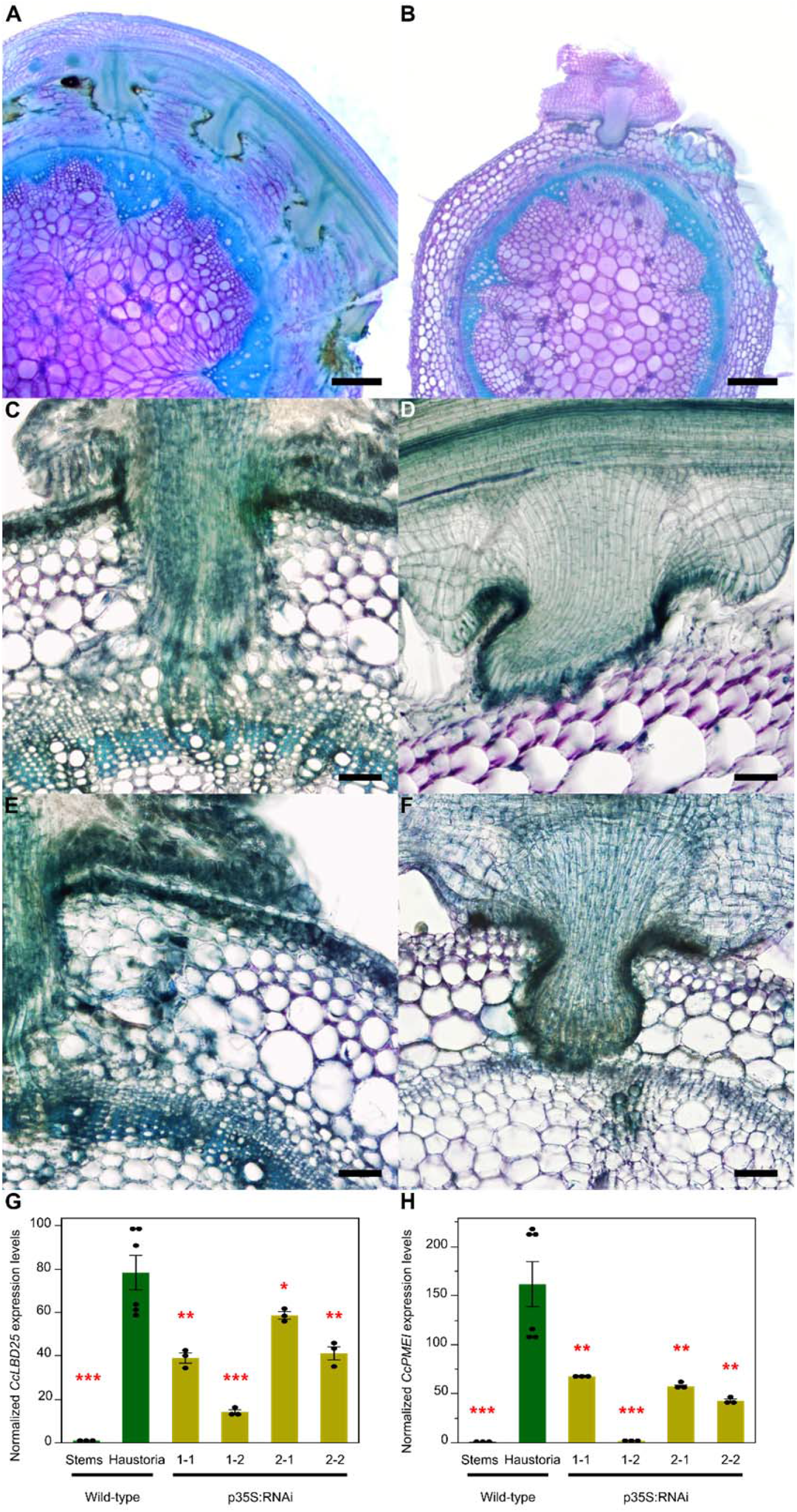
Haustorium phenotypes and gene expression levels of *C. campestris* growing on HIGS *CcLBD25* RNAi transgenic plants. (A, C, E) *C. campestris* haustoria growing on a wild-type H1706 host. (B, D, F) *C. campestris* haustoria growing on a *CcLBD25* RNAi transgenic tomato. (A, B) Scale bars = 500 µm. (C, D, E, F) Scale bars = 100 µm. (G, H) Expression levels of *CcLBD25* and *CcPMEI* in *C. campestris* haustoria grown on wild-type tomatoes and *CcLBD25* RNAi transgenic plants. Data presented are assessed using one-tailed Welch’s t-test with wild-type haustoria as control. “*” p-value < 0.05. “**” p-value < 0.005. “***” p-value < 0.001. p35S:RNAi indicates the transgenic plants with the 35S promoter driving *CcLBD25* RNAi construct.

We also noticed the down-regulation of *CcLBD25* influenced the parasite penetration process. Fresh tissue sections of the haustoria growing on wild types showed a clear zone in tomato cortex tissues near haustorium tissues (Fig. 6A, C, E). Since the metachromatic staining of Toluidine Blue O is based on cell wall composition and pH values and a pink to purple color indicates pectin presence, this result indicates that the pectins in tomato cortex tissues may have been digested or the pH condition in the cell wall has been changed in the haustorium penetrating process (Fig. 6A, C, E). On the other hand, the *C. campestris* growing on *CcLBD25* RNAi transgenic tomatoes still showed pink to purple color in the cortex near the haustorium attachment sites (Fig. 6B, D). Hence, less pectin digestion or cell wall modification happened in tomato cortex tissues near these *CcLBD25* downregulated haustorium tissues compared to the haustoria growing on wild-type. These haustorium structural phenotypes correlate well with our SOM9 GCN from the *C. campestris* tissue transcriptome. *CcLBD25* is one of the transcription factor central hub genes in module 1 with many first and second layers of connection with genes involved in cell wall modification, including pectin lyases and pectin methyl-esterase inhibitors (PMEIs). Based on many previous studies, the interplay between pectin methylesterase (PME) and PMEI is an important determinant of cell wall loosening, strengthening, and organ formation (Wormit and Usadel, 2018). Therefore, we hypothesized that PMEIs might be one of the key regulators that cause the haustorium phenotype in *CcLBD25* downregulated haustorium tissues. To test if the down-regulation of *CcLBD25* would affect *PMEI* expression levels, we conducted qPCR to detect *CcPMEI* expression levels in the tissues of *C. campestris* plants that are growing on *CcLBD25* RNAi transgenic plants. Our results show that *CcPMEI* expression levels are also significantly reduced when *CcLBD25* is knocked-down (Fig. 6G, H). Thus *CcLBD25* might directly or indirectly regulate *CcPMEI* at the transcriptional level. These results verify the hypothesis that *CcLBD25* plays an important role in haustorium development and might regulate cell wall modification.

### Investigating the impact of *CcLBD25* on early-stage haustorium development using an *in vitro* haustoria (IVH) system

Previous studies indicate that several auxin-inducible *LBD* genes function in lateral root initiation (Goh et al., 2012). We noticed that auxin efflux carriers and auxin-responsive genes are also in the SOM9 gene co-expression network (Fig. 2). Therefore, we proposed that *CcLBD25* might regulate early-stage haustorium development in *C. campestris*. In order to assay the role of *CcLBD25* in *C. campestris* haustorium initiation, we developed an *<u>i</u>n <u>v</u>itro* <u>h</u>austorium (IVH) system coupled with HIGS (Fig. 7A). This method is inspired by the previous discovery that *Cuscuta* haustoria can be induced by physical contact and far-red light signals (Tada et al., 1996) and many studies confirmed that small RNAs and mRNAs can move cross species through haustorial phloem connection (David-Schwartz et al., 2008; Alakonya et al., 2012; Kim et al., 2014; Johnson et al., 2019). Therefore, we took the *C. campestris* strands near the haustorium attachment sites growing on wild-type and *CcLBD25 RNAi* transgenic tomato (Fig. 7B) and sandwiched these strands in between two layers of agar to provide sufficient physical contact signals (Fig. 7A, C). We then illuminated these plates under far-light for 5 days at which point prehaustoria are readily visible (Fig. 7D, E). Since the IVH induction is rapid and these prehaustoria can be easily separated from the agar, this method allowed us to count prehaustoria numbers under the microscope and validate the effect of *CcLBD25* RNAi on haustorium initiation. The strands from the *C. campestris* grown on *CcLBD25 RNAi* transgenic tomatoes produced many fewer prehaustoria than the strands from those grown on wild types (Fig. 7F). This result indicates that reduced *CcLBD25* expression impeded haustorium initiation and confirms that *CcLBD25* is a key regulator of early-stage haustorium development, as suggested by our LCM RNA-Seq analysis results.

**Figure 7.**
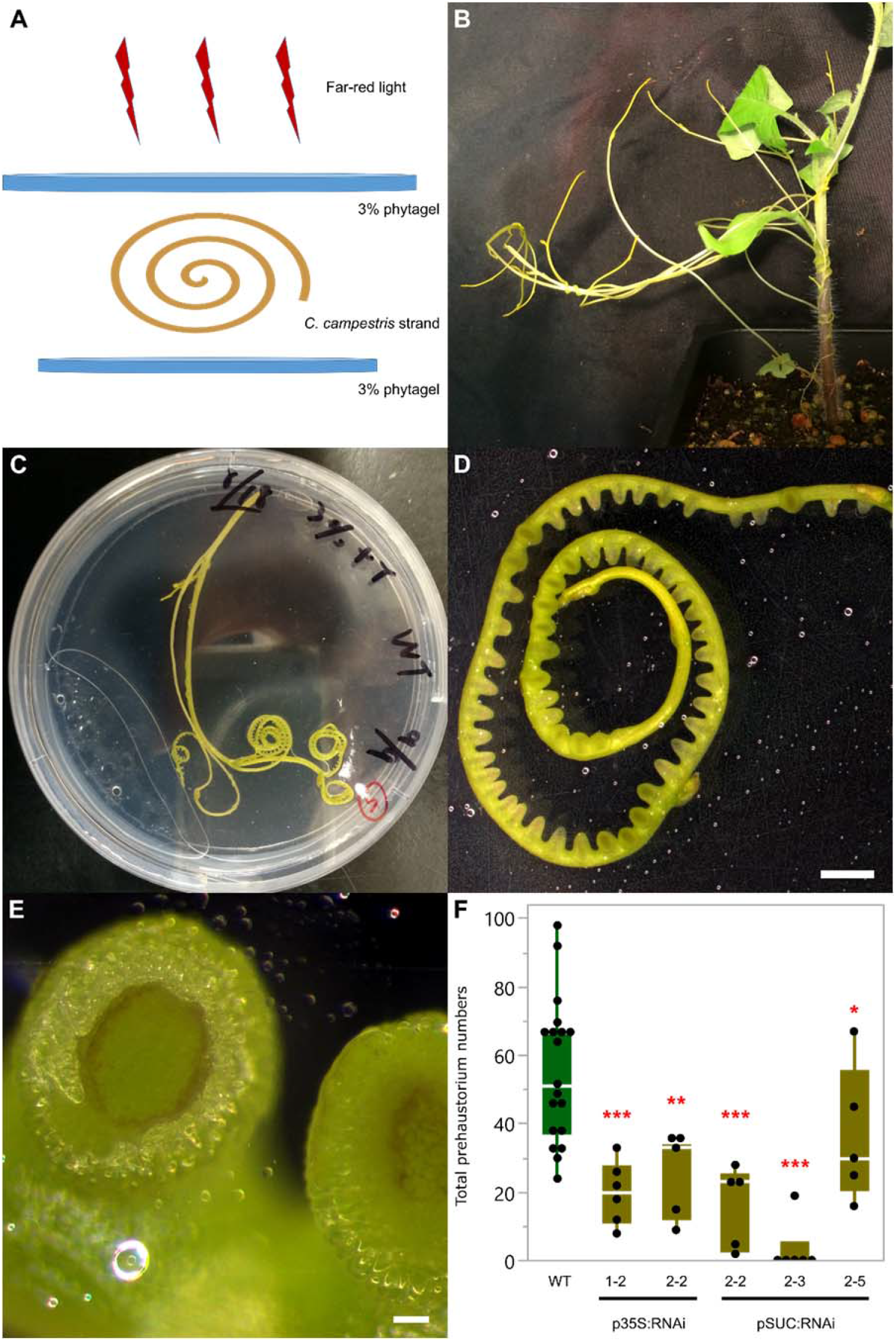
Far-red light-induced *in vitro* haustorium (IVH) phenotypes of *C. campestris* growing on HIGS *CcLBD25* RNAi transgenic plants. (A) An illustration of the setup for the IVH system. (B) *C. campestris* strands near the haustorium attachment sites. (C) An IVH plate with a *C. campestris* strand sandwiched in between two layers of agar to provide sufficient physical contact signals. (D, E) After illuminating these plates under far-light for 5 days, prehaustoria are readily visible. (D) Scale bar = 2 mm. (E) Scale bars = 100 µm. (F) *C. campestris* strands were detached and subjected to IVH and the number of prehaustoria were counted. Data presented are assessed using one-tailed Welch’s t-test with wild-type (WT) as control. “*” p-value < 0.06. “**” p-value < 0.001. “***” p-value < 0.0005. p35S:RNAi indicates the transgenic plants with the 35S promoter driving *CcLBD25* RNAi construct. pSUC:RNAi indicates the transgenic plants with the SUC2 promoter driving *CcLBD25* RNAi construct.

## Discussion

In this study, we demonstrate that *CcLBD25* is a crucial regulator of several aspects of *C. campestris* haustorium development, including haustorium initiation, cell-wall loosening, and searching hyphae growth. We use transcriptome of six *C. campestris* tissue types and RNA-Seq data of LCM captured haustoria at three developmental stages to reveal the potential molecular mechanisms and the complexity of gene networks during the haustorium formation process. Our results provide a more comprehensive analysis of the *CcLBD25* centered regulatory system and illustrate that *CcLBD25* might directly or indirectly coordinate different groups of genes that are expressed only at the early or mature stage during haustorium development.

### Lateral root development and haustorium development

In non-parasitic plants, like Arabidopsis, *AtLBD25* was also named *DOWN IN DARK AND AUXIN1* (*DDA1*) because *lbd25* mutant plants showed reduced sensitivity to auxin and reduced number of lateral roots (Mangeon et al., 2010). These phenotypes indicate that *AtLBD25* functions in lateral root formation by promoting auxin signaling (Mangeon et al., 2010). In the root parasitic plant, *Thesium chinense, TcLBD25* was highly expressed during haustorium formation (Ichihashi et al., 2017). This supports the hypothesis that root parasitic plants co-opted the lateral root formation machinery into haustorium organogenesis. However, whether rootless stem parasitic plants *Cuscuta* spp. also followed the same path to generating haustoria was unknown. In this study, we identified *CcLBD25* as playing a key role in *Cuscuta* haustorium development. Our SOM9 gene co-expression network shows that auxin efflux carriers and auxin-responsive genes are also connected with *CcLBD25* (Fig. 2). These pieces of evidence suggest that *Cuscuta* spp. adopted not only the shoot developmental programs (Alakonya et al., 2012) but also the lateral root programming system into haustorium organogenesis.

### Searching hyphae development

Down-regulating *CcLBD25* reduced searching hyphae formation (Fig. 6B, D, F), indicating that *CcLBD25* is involved in searching hyphae development. Surprisingly, *AtLBD25* is not only expressed in roots but also expressed in pollen (Mangeon et al., 2010). Previous reports indicate that *AtLBD25* is especially highly expressed during the pollen late developmental stage (Kim et al., 2016). Intriguingly, many genes that are involved in haustoria development also play important roles in flower and pollen development (Yang et al., 2015; Yoshida et al., 2019). Recent research on haustoria 3D structure also indicates that the growth pattern of intrusive cells is similar to the rapid polar growth of pollen tubes (Masumoto et al., 2020). Taken together with our results in this study and previous findings in other organisms, we suggest that the genes that are regulating pollen development or pollen tube growth, like *LBD25*, might be adopted by parasitic plants for haustorium intrusive cell and searching hyphae development. This discovery also confirmed the hypothesis that parasitic plants co-opted the developmental reprogramming process from multiple sources instead of just a single organ.

### Cell adhesion and cell wall loosening in parasitism

The mechanical properties and chemical conditions of cell walls have been reported to be critical for regulating plant organ morphogenesis (Chebli and Geitmann, 2017; Zhao et al., 2018). By remodeling cell wall composition or extracellular environments, plants generate local cell wall loosening and strengthening, which allows anisotropic growth processes to occur (Chebli and Geitmann, 2017). Recent studies also indicate that the interaction between pectin and other cell wall components is an important determinant for plant organogenesis (Chebli and Geitmann, 2017; Saffer, 2018) and the interplay between PME and PMEI plays a vital role in regulating physical properties of the cell wall (Wormit and Usadel, 2018). In the root parasitic plant, *Orobanche cumana*, a PME is shown to be present at the host and parasite interface and have pectolytic activity (Losner-Goshen et al., 1998). These results suggest that parasitic plants produce PME to degrade pectin in the host cell wall and help with haustorium penetration. Our SOM9 GCNs shows that *CcLBD25* is co-expressed with many pectin lyases and PMEIs (Fig. 2B, 3C, 3D, 4C, 4D), implying that *CcLBD25* might be the key transcription factor regulating expression of the enzymes involved in pectin remodeling. The haustoria grown on *CcLBD25* RNAi transgenic plants failed to penetrate host tissues and were unable to create a clear zone at the host and parasite interface (Fig. 6A-F), supporting the tight connection between *CcLBD25* and pectin-modifying enzymes. *CcLBD25* and PMEIs co-expressed in the mature stage of haustorium (Fig. 4C-D), when cell wall loosening occurs for haustorium penetration.

On the other hand, since the patterns of de-methylesterification on homogalacturonans (HG) determines cell wall loosening or strengthening, pectin properties also play a role in cell adhesion, which is regulated by PME and PMEI (Wormit and Usadel, 2018). Previous studies also indicate that *Cuscuta* spp. secrete pectin-rich adhesive materials to help with adhesion and allow attachment to their hosts (Vaughn, 2002; Shimizu and Aoki, 2019). This is consistent with our discovery that both *CcLBD25* and *PMEI*s are highly expressed in the early stage of haustorium development, which would be responsible for the adhesion process in *C. campestris* (Fig. 4C-D).

## Conclusions

Our detailed bioinformatic analysis on previously published *C. campestris* tissue type transcriptome coupled with LCM of RNA-Seq data from three haustorium developmental stages helped us hone in on the molecular mechanism of parasitic plant haustorium development. The discovery that *CcLBD25* plays a pivotal role in many aspects of haustorium formation shows that the regulatory machinery of haustorium development is potentially shared by both root and stem parasites. Although previous studies have indicated that parasitic plants evolved independently in about 13 different families, this conserved molecular mechanism supports the hypothesis that stem parasitic plants also adopted the programming of lateral root formation in non-parasitic plants into haustorium development. The results of this study not only provide an insight into molecular mechanisms by which LBD25 may regulate parasitic plant haustorium development but also raise potential for developing a universal parasitic weed-resistant crop that can defend both stem and root parasitic plants at the same time.

## Materials and Methods

### *Cuscuta campestris* materials

We thank W. Thomas Lanini for providing dodder seeds collected from tomato field in California. These dodder materials were previously identified as *Cuscuta pentagona* (Yaakov et al., 2001), a closely related species to *Cuscuta campestris (Costea et al*., *2015)*. We use molecular phylogenetics of plastid *trnL-F* intron / spacer region, plastid ribulose-1,5-bisphosphate carboxylase/oxygenase large subunit (*rbcL*), nuclear internal transcribed spacer (*nrITS*), and nuclear large-subunit ribosomal DNA (*nrLSU*) sequences (Stefanović et al., 2007; García et al., 2014; Costea et al., 2015) to verify our dodder isolate is the same as *Cuscuta campestris* 201, Rose 46281 (WTU) from USA, CA (Jhu et al., 2020) by comparing with published sequences (Costea et al., 2015).

### RNA-Seq data mapping and processing

For *C. campestris* tissue type RNA-Seq analysis, we used the raw data previously published (Ranjan et al., 2014). This RNA-Seq data contain six different *C. campestris* tissues, including seeds, seedlings, stems, prehaustoria, haustoria, and flowers, grown on the tomato (*Solanum lycoperscum*) Heinz 1706 (H1706) cultivar and *Nicotiana benthamiana* (*N. benthamiana*). We mapped both *C. campestris* tissue type and LCM RNA-Seq data to the genome of *C. campestris* (Vogel et al., 2018) with Bowtie 2 (Langmead and Salzberg, 2012) and used EdgeR (Robinson et al., 2009) to get normalized trimmed mean of M values (TMM) for further analysis.

### MDS and PCA with SOM Clustering

After normalization steps, we used cmdscale in R stats package to create multidimensional scaling (MDS) data matrix and then generate MDS plots. For *C. campestris* tissue types RNA-Seq data, we selected genes with coefficient of variation > 0.85 for PCA analysis. We calculated principal component values using prcomp function in R stats package. Selected genes are clustered for multilevel six-by-two hexagonal SOM using som function in the kohonen package (Wehrens and Buydens, 2007). We visualized the SOM clustering results in PCA plots. For *C. campestris* LCM RNA-Seq data, genes in the upper 50% quartile of coefficient of variation were selected for further analysis. Selected genes were then clustered for multilevel three-by-two hexagonal SOM.

### Construct Gene coexpression networks

We use the genes that are classified in selected SOM groups to build GCNs. The R script is modified from our previously published method (Ichihashi et al., 2014) and the updated script is uploaded to GitHub and included in code availability. The SOM9 GCNs for *C. campestris* tissue type data was constructed with normal quantile cutoff = 0.93. The SOM2+3+9 GCNs for *C. campestris* tissue type data was constructed with normal quantile cutoff =0.94. For the GCN of *C. campestris* LCM data, we used the SOM9 gene list from tissue type RNA-Seq and construct the GCN of these genes based on the expression profiles in LCM RNA-Seq data with normal quantile cutoff =0.94. These networks were then visulaized using Cytoscape version 3.8.0.

### Functional annotation and GO enrichment analysis of RNA-Seq data

Since many genes are not functionally annotated in the recently published *C. campestris* genome (Vogel et al., 2018), we used BLASTN with 1e-5 as an e-value threshold to compare our previously annotated transcriptome final contigs with current *C. campestris* genome genes and only keep the top 1 scored hits for each gene (Supplementary Table 1). After we obtain this master list, we combined the functional annotation of our published transcriptome based on NCBI nonredundant database and TAIR10 (Ranjan et al., 2014) with *C. campestris* genome gene IDs to create a more complete functional annotation (Supplementary Table 1). TAIR ID hits are used for GO Enrichment Analysis on http://geneontology.org/ for gene clusters and modules.

### LCM RNA-seq Library Preparation and Sequencing

We infested about four-leaves-stage Heinz 1706 tomato plants with *C. campestris* strands. Tomato stems with haustoria are collected at 4 days post attachment (DPA) and fixed in formaldehyde – acetic acid – alcohol (FAA). These samples were dehydrated by the ethanol series and embedded in paraffin (Paraplast X-TRA, Thermo Fisher Scientific). We prepared 10 μm thick sections on a Leica RM2125RT rotary microtome. Tissue was processed within one month of fixation to ensure RNA quality. Haustorial tissues of the 3 defined developmental stages were dissected on a Leica LMD6000 Laser Microdissection System. Tissue was collected in lysis buffer from RNAqueous-Micro Total RNA Isolation Kit (Ambion) and stored at −80 **°**C. RNA was extracted using RNAqueous-Micro Total RNA Isolation Kit (Ambion) and amplified using WT-Ovation Pico RNA Amplification System (ver. 1.0, NuGEN Technologies Inc.) following manufacturer instructions. RNA-seq libraries for Illumina sequencing were constructed following a previously published method (Kumar et al., 2012) with slight modifications. Libraries were quantified, pooled to equal amounts, and their quality was checked on a Bioanalyzer 2100 (Agilent). Libraries were sequenced on a HiSeq2000 Illumina Sequencer at the Vincent J Coates Genomics Sequencing Laboratory at UC Berkeley.

### *CcLBD25* RNAi transgenic plants and HIGS efficiency verification

We used pTKO2 vector (Snowden et al., 2005; Brendolise et al., 2017), which enables streamlined cloning by using two GATEWAY cassettes, positioned at opposite directions, separated by an Arabidopsis ACT2 intron and under the control of the 35S constitutive promoter. We have previously shown that producing the RNAi construct at phloem cells specifically using the SUC2 promoter was effective at dodder HIGS (Alakonya et al., 2012). Therefore, we replaced the 35S promoter with the SUC2 promoter and generated pTKOS (Supplemental Fig. 8). We used BLAST to identify a 292 bp fragment that was specific to *CcLBD25* and different from tomato genes. This RNAi fragment was amplified from *C. campestris* gDNA, TOPO cloned into pCR8/GW-TOPO (Life Technologies) and LR recombined into pTKO2 and pTKOS. These constructs were then sent to the UC Davis Plant Transformation Facility to generate *CcLBD25* RNAi transgenic tomato plants.

All T0 transgenic plants were selected by kanamycin resistance and their gDNAs were extracted and PCR performed to verify they have *CcLBD25* RNAi constructs. To validate HIGS efficiency and quantify the expression level of *CcLBD25* and *CcPMEI* in *C. campestris*, dodder tissues were harvested from both *C. campestris* grown on wild-type plants and T2 *CcLBD25* RNAi transgenic plants. We froze tissues in liquid nitrogen and ground them in extraction buffer using a bead beater (Mini Beadbeater 96; BioSpec Products). Following our previously published poly-A based RNA extraction method (Townsley et al., 2015), we obtained total mRNA from *C. campestris* and then used Superscript III reverse transcriptase (Invitrogen) for reverse transcription to synthesize cDNA as described by the manufacturer instructions. Real-time qPCR was performed using a Bio-Rad iCycler iQ real-time thermal cycler with Bio-Rad IQ SYBR Green super mix.

### Whole-plant phenotype assays

Based on previous studies, many plant species are reported to have early flowering phenotypes in response to environmental stresses (Wada and Takeno, 2010; Riboni et al., 2014). Therefore, we grew *C. campestris* on wild-type Heinz 1706 tomatoes and *CcLBD25* RNAi T1 transgenic tomato plants and then quantified how fast these *C. campestris* plants transition to their reproductive stage. The number of *C. campestris* plants that transitioned to the flowering stage were counted at 9, 10, 14 days post attachment (DPA) to test whether a stress-induced flowering phenotype could be observed.

To quantify the effect of *CcLBD25* downregulation on *C. campestris* growth, we infested 3-weeks-old tomato plants with about 10 cm stem segments *C. campestris*, which originally are grown on wild-type H1706. We harvested all *C. campestris* tissues grown on wild-type H1706 and *CcLBD25* RNAi T2 transgenic plants at 14 DPA. These *C. campestris* tissues were then carefully separated from their host plant stems by hands and their fresh weights were measured using chemical weighing scales.

### *in vitro* haustoria (IVH) system

Inspired by the previous discovery that *Cuscuta* haustoria can be induced by physical contact and far-red light signals (Tada et al., 1996), we developed an *in vitro* haustoria (IVH) system for haustorium induction without hosts. In this method, we detached *Cuscuta* stem segments that are right next to a stable haustorium attachment from the *C. campestris* grown on wild-type plants and T2 *CcLBD25* RNAi transgenic plants. *Cuscuta* strands with shoot apices detached from a host plant are sandwiched between 3% Phytagel agar containing 0.5X Murashige and Skoog medium to provide tactile stimuli (Fig. 7A-C). These combined plates then irradiated with far-red light for two hours. After 5 days of growth in darkness in a 22 °C growth chamber, prehaustoria are readily visible (Fig. 7D, E). We then counted the number of prehaustoria under a Zeiss SteREO Discovery, V12 microscope for quantification. Since the RNAi silencing signal is systemic (Alakonya et al., 2012; David-Schwartz et al., 2008) and IVH induction is rapid, we can validate the effect of *CcLBD25* RNAi on haustoria development.

### Fresh tissue sectioning and histology

For fresh vibratome sections of haustoria attached to wild-type and *CcLBD25 RNAi* host stems, we collected samples and embedded them in 7% Plant Tissue Culture Agar. We then fixed these agar blocks in FAA (final concentration: 4% formaldehyde, 5% glacial acetic acid, and 50% ethanol) overnight, 50% ethanol for one hour, and then transferred to 70% ethanol for storage. These agar blocks were then sectioned using Lancer Vibratome Series 1000 to prepare 100 μm sections. We kept these sections in 4°C water and then conducted Toluidine Blue O Staining. We followed the published protocol (O’Brien et al., 1964) with some modifications. The sections were immersed in the stain for 30 seconds, and then washed them with water three times for 30 seconds each. After removing the agar from around the sections using forceps, we mounted the sections with water on a slide and imaged using a Zeiss SteREO Discovery, V12 microscope, and a Nikon Eclipse E600 microscope.

## Acknowledgements

We are grateful to the UC Davis Plant Transformation Facility for generating *CcLBD25* RNAi transgenic tomato plants. We thank Kristina Zumstein for help in maintaining transgenic tomato seed stocks, and Richard Philbrook, Kaiwen Zhang, and Junqi Lu for helping with some parts of experiments and vibratome sectioning. We also thank Aaron Leichty, Steven Rowland and Karo Czarnecki for their input on bioinformatics analyses.

## Funding

This work was funded by USDA-NIFA (2013-02345). M.-Y. J. was supported by Yen Chuang Taiwan Fellowship, Taiwan Government Scholarship (GSSA), Elsie Taylor Stocking Memorial Fellowship, Katherine Esau Summer Graduate Fellowship, Loomis Robert S. and Lois Ann Graduate Fellowship in Agronomy, and the UCD Graduate Research Award. Y. I. was supported by Grant-in-Aid for Young Scientists from the Ministry of Education, Culture, Sports, Science and Technology, Japan [B; grant no. 15K18589 to YI].

## Competing interests

The authors declare that they have no competing interests.

## Code availability

Updated R scripts for MDS, PCA and SOM analysis and gene coexpression network analysis are all deposited on GitHub (Link: https://github.com/MinYaoJhu/CcLBD25_project.git).

## Data availability

All data is available in the main text or the supplementary materials. LCM RNA-Seq raw data are deposited on NCBI SRA PRJNA687611.

